# Discovery of baicalin and baicalein as novel, natural product inhibitors of SARS-CoV-2 3CL protease *in vitro*

**DOI:** 10.1101/2020.04.13.038687

**Authors:** Haixia Su, Sheng Yao, Wenfeng Zhao, Minjun Li, Jia Liu, WeiJuan Shang, Hang Xie, Changqiang Ke, Meina Gao, Kunqian Yu, Hong Liu, Jingshan Shen, Wei Tang, Leike Zhang, Jianping Zuo, Hualiang Jiang, Fang Bai, Yan Wu, Yang Ye, Yechun Xu

## Abstract

Human infections with severe acute respiratory syndrome coronavirus 2 (SARS-CoV-2) cause coronavirus disease 19 (COVID-19) and there is currently no cure. The 3C-like protease (3CLpro), a highly conserved protease indispensable for replication of coronaviruses, is a promising target for development of broad-spectrum antiviral drugs. To advance the speed of drug discovery and development, we investigated the inhibition of SARS-CoV-2 3CLpro by natural products derived from Chinese traditional medicines. Baicalin and baicalein were identified as the first non-covalent, non-peptidomimetic inhibitors of SARS-CoV-2 3CLpro and exhibited potent antiviral activities in a cell-based system. Remarkably, the binding mode of baicalein with SARS-CoV-2 3CLpro determined by X-ray protein crystallography is distinctly different from those of known inhibitors. Baicalein is perfectly ensconced in the core of the substrate-binding pocket by interacting with two catalytic residues, the crucial S1/S2 subsites and the oxyanion loop, acting as a “shield” in front of the catalytic dyad to prevent the peptide substrate approaching the active site. The simple chemical structure, unique mode of action, and potent antiviral activities *in vitro*, coupled with the favorable safety data from clinical trials, emphasize that baicalein provides a great opportunity for the development of critically needed anti-coronaviral drugs.

---

Coronaviruses (CoVs; subfamily Coronavirinae, family Coronaviridae, order Nidovirales), are a group of highly diverse, enveloped, positive-sense, single-stranded RNA viruses which are prevalent pathogens responsible for a wide range of diseases in animals as well as humans (*1*, *2*). Human CoV strains including 229E, NL63, OC43, and HKU1 typically lead to mild to moderate upper respiratory tract infections such as colds and pneumonia, whereas human infections with zoonotic CoVs, such as severe acute respiratory syndrome (SARS) in 2003 and Middle East respiratory syndrome (MERS) in 2012, caused severe respiratory diseases with high morbidity and mortality. SARS-CoV-2 is a new human coronavirus identified recently and appears to be more readily transmitted from human to human (*3*, *4*). It is 96% identical at the whole genome level to a bat coronavirus and shares 79.5% sequence identify to SARS-CoV (*5*). Human infections with SARS-CoV-2 cause severe and fatal pneumonia. As of April 7, 2020, more than one million cases have been diagnosed and approximately 80000 patients have died of coronavirus disease 19 (COVID-19) worldwide, posing a significant threat to public health globally. It also marks the third introduction of a highly pathogenic CoV into the human population in the 21st century. Unfortunately, no efficacious antiviral drug and vaccine have been approved for the prophylaxis or treatment of the highly virulent SARS-CoV, MERS-CoV, or SARS-CoV-2 infections in humans.

Featuring the largest genomes among the RNA viruses, CoVs share key genomic elements that provide promising therapeutic targets (*1*, *6*). A chymotrypsin-like cysteine protease called 3C-like protease (3CLpro, 11 cleavage sites in the polyproteins) together with a papain-like protease (PLpro, three cleavage sites in the polyproteins) is required to process polyproteins into mature non-structural proteins such as RNA-dependent RNA polymerase and helicase, which are essential for viral transcription and replication. Moreover, the substrate specificity of the 3CLpro, also known as the main protease (Mpro), is highly conserved among different CoVs and is similar to that of the main picornavirus 3C protease, thereby making it an ideal target for the development of broad-spectrum antiviral drugs (*6*, *7*). Although several types of substrate-like peptidomimetic inhibitors of SARS-CoV 3CLpro or MERS-CoV 3CLpro, mostly with a mode of covalent action, have been reported, none has yet progressed into clinical trials (*1*, *6*). Classical small molecules with more-drug-like properties and their binding with 3CLpros supported by complex structure determination have not yet emerged.

Traditional Chinese medicines (TCMs) have evolved over thousands of years and are an invaluable source for drug discovery and development. As a notable example, the discovery of artemisinin (Qinghaosu), which was originally isolated from the TCM *Artemisia annua* L. (Qinghao), is a milestone in the treatment of malaria. TCMs as well as purified natural products also provide a rich resource for novel antiviral drug development. Several herbal medicines and natural products have shown antiviral activities against viral pathogens (*8*–*11*). Among these, the roots of *Scutellaria baicalensis* Georgi (Huangqin in Chinese) are frequently used in TCM for the prophylaxis and treatment of hepatitis and respiratory disorders (*12*–*14*). In the present study, an enzymatic assay was performed to test if the ingredients isolated from *S. baicalensis* are inhibitors of SARS-CoV-2 3CLpro. As a result, baicalin and baicalein, two bioactive components from *S. baicalensis*, are identified as novel inhibitors of SARS-CoV-2 3CLpro with an antiviral activity in the SARS-CoV-2 infected cells. A crystal structure of SARS-CoV-2 3CLpro in complex with baicalein, the first non-covalent, non-peptidomimetic small-molecule inhibitor, was also determined, revealing a unique binding mode of this natural product with the protease.

## Inhibition of SARS-CoV-2 3CLpro by baicalin and baicalein

The pivotal role of the 3CL protease in processing polyproteins into individual functional proteins for viral replication and a highly conserved substrate specificity of the enzyme among various CoVs make it a promising target for screening of inhibitors. A fluorescence resonance energy transfer (FRET) protease assay was applied to measure the proteolytic activity of the recombinant SARS-CoV-2 3CLpro on a fluorogenic substrate. The detail of the assay is described in the Experimental section. This FRET-based protease assay was utilized to screen natural products as novel inhibitors of SARS-CoV-2 3CLpro. It was first used to determine the inhibitory activities of the total aqueous extract, fractionations, and purified compounds from *S. baicalensis* against SARS-CoV-2 3CLpro (see Supplementary Materials, Fig. S1). As the result, two fractions from *S. baicalensis* showed significant inhibition on SARS-CoV-2 3CLpro at 10.0 μg/mL (Table S1). Surprisingly, baicalin, the major component in fraction 8, shows an IC_50_ of 6.41 μM against the protease, while baicalein, the major component in fraction 12, has an IC_50_ of 0.94 μM (Fig. S2; Table 1). Accordingly, baicalin and baicalein are identified as novel non-peptidomimetic inhibitors of SARS-CoV-2 3CLpro with single-digit micromolar potency.

**Table 1.** Inhibitory activity and binding affinity of baicalin and baicalein with SARS-CoV-2 3CLpro determined by enzymatic assay, ITC^1^, and ESI-MS^2^.

| Compounds | $K_d$ (μM)<sup>ITC</sup> | $\Delta G$ (kJ/mol) | $\Delta H$ (kJ/mol) | $-T\Delta S$ (kJ/mol) | $K_d$ (μM)<sup>MRMS</sup> | IC<sub>50</sub> (μM) |
| --- | --- | --- | --- | --- | --- | --- |
| Baicalin | 11.50 ± 0.37 | -28.21 ± 0.42 | -2.77 ± 0.51 | -25.44 ± 0.89 | 12.73 ± 0.81 | 6.41 ± 0.95 |
| Baicalein | 4.03 ± 0.37 | -30.78 ± 0.24 | -5.64 ± 1.31 | -25.14 ± 1.09 | 1.40 ± 0.10 | 0.94 ± 0.20 |

<sup>1</sup>ITC means isothermal titration calorimetry.
<sup>2</sup>ESI-MS means Native state electrospray ionization mass spectrometry.

## Protein-ligand binding supported by ITC and ESI-MS

To validate the binding of baicalin and baicalein with SARS-CoV-2 3CLpro and exclude the suspicion of being the pan-assay interference compounds (PAINS) (*15*), their binding affinities with the protease were measured by isothermal titration calorimetry (ITC), widely known as an invaluable tool used to determine thermodynamic parameters of protein-ligand interactions such as *K*_d_ (Fig. 1, A and B; Table 1). The resulting *K*_d_ of baicalin and baicalein binding with SARS-CoV-2 3CLpro is 11.50 and 4.03 μM, respectively, which has a good correlation with the IC_50_s mentioned above, demonstrating that specific binding of the compounds with the enzyme is responsible for their bioactivities. Moreover, the ITC profiles in combination with their chemical structures suggest that baicalin and baicalein act as non-covalent inhibitors of SARS-CoV-2 3CLpro with a high ligand binding efficiency.

**Fig. 1.**
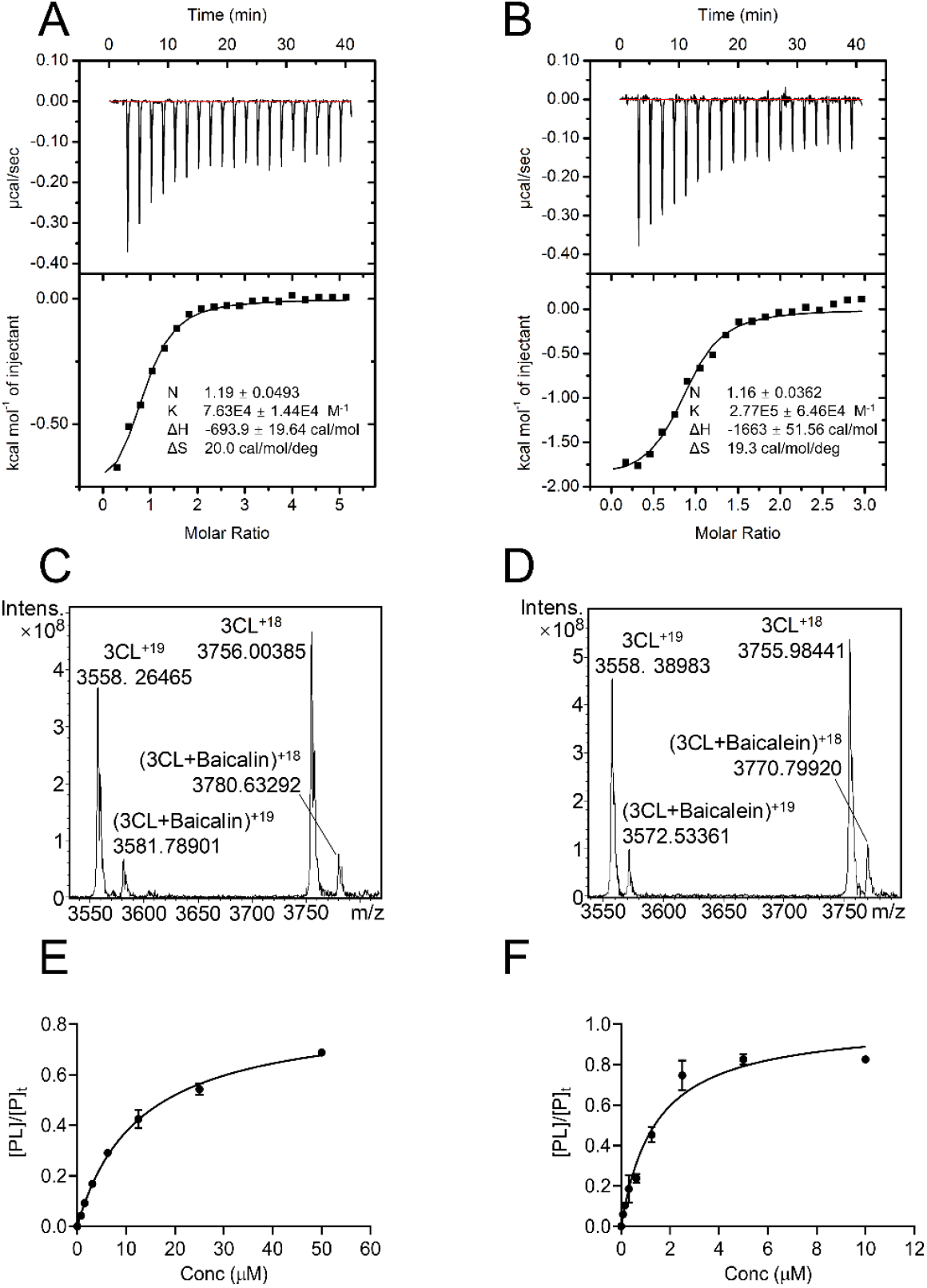
Binding of baicalin and baicalein with SARS-CoV-2 3CLpro characterized by isothermal titration calorimetry (ITC) and native state mass spectrometry. (**A** and **B**) Representative thermodynamic profiles of baicalin (A) and baicalein (B) binding with the protease in solution resulted from ITC measurements. (**C** and **D**) Native state mass spectrometry spectra of SARS-CoV-2 3CLpro with baicalin and baicalein: (C) 1-μM SARS-CoV-2 3CLpro with 3.13-μM baicalin; (D) 1-μM SARS-CoV-2 3CLpro with 0.31-μM baicalein. 3CL in two plots represents SARS-CoV-2 3CLpro. (**E** and **F**) Binding affinities of baicalin (E) and baicalein (F) with SARS-CoV-2 3CLpro determined by native state mass spectrometry. Fraction of ligand bound protein ([PL]/P*100) obtained under optimized conditions plotted against total ligand concentrations.

Native state electrospray ionization mass spectrometry (ESI-MS) has been used extensively to directly observe native state proteins and protein complexes, allowing direct detection of protein-ligand non-covalent complexes with *K*_d_s as weak as 1 mM (*16*). The determination of Δ*m*/*z* between [protein + ligand] *m/z* and [unbound protein] *m/z* is able to identify a ligand as a binder with the correct molecular weight, while the ratio of the intensity of the [protein + ligand] peaks relative to [unbound protein] peaks provides a qualitative indication of the ligand-binding affinity. Herein, an ESI-MS analysis using high-resolution magnetic resonance mass spectrometry (MRMS) was carried out to detect the binding of baicalin and baicalein with SARS-CoV-2 3CLpro. For the free protease performance optimization, the mass range around the change stated 18+ was isolated with a center mass of the quadrupole of *m/z* 3750 (Fig. S3). For the ligand-binding screening studies, two charge states (18+ and 19+) have been used for calculation of the free protease and protein-ligand complex intensities. The representative spectra of samples containing SARS-CoV-2 3CLpro (1 μM) and baicalin (3.13 μM) or baicalein (0.31 μM) acquired under both optimized and screening conditions are shown in Fig. 2C and D, demonstrating a specific binding of baicalin or baicalein with the protease. Moreover, the plot of the fraction of the bound protease versus the total concentration of baicalin or baicalein obtained *K*_d_s of 12.73 and 1.40 μM for baicalin and baicalein, respectively (Fig. 1E and F), in keeping with the results from the ITC measurements.

**Fig. 2.**
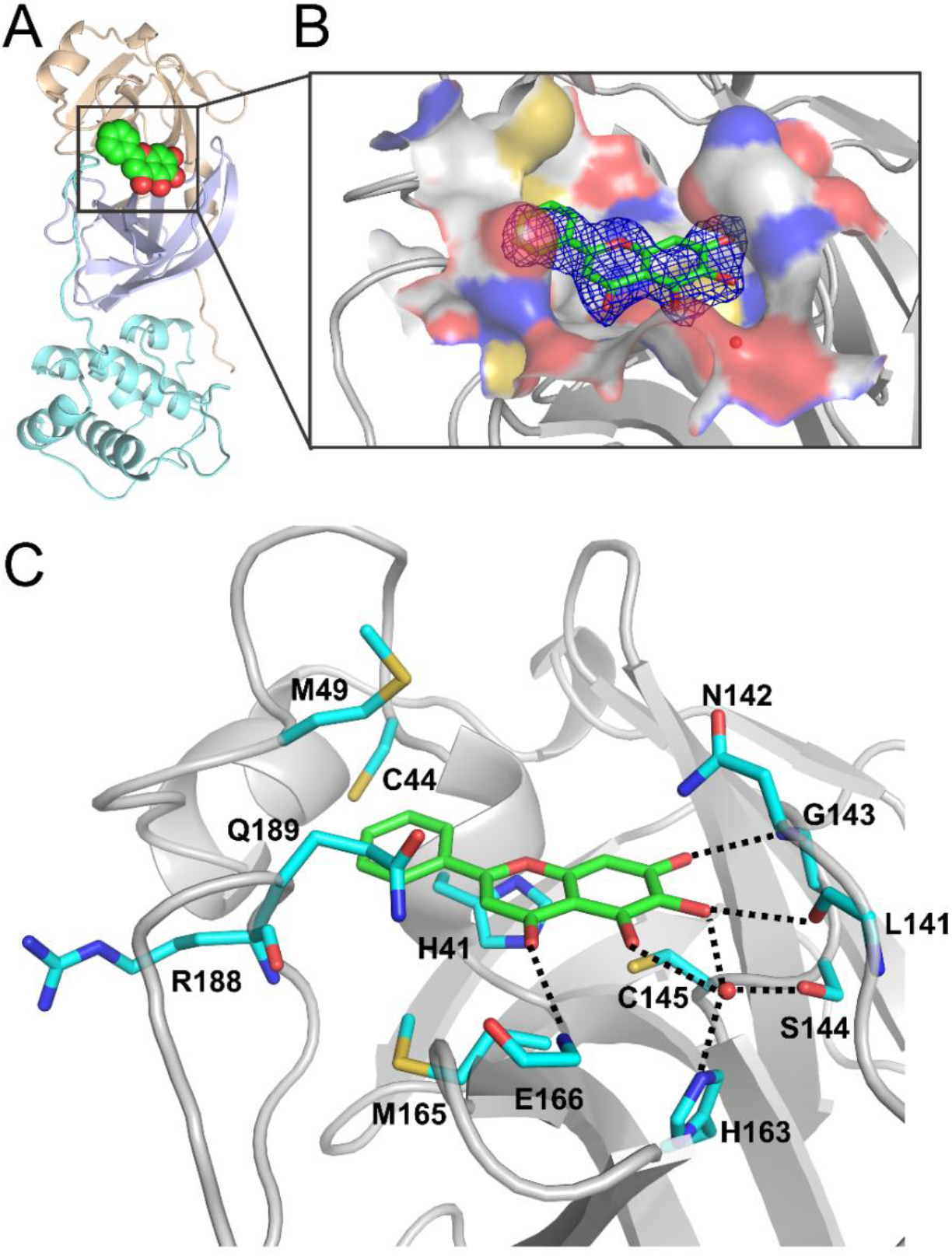
Crystal structure of SARS-CoV-2 3CLpro in complex with baicalein (PDB code 6M2N). (**A**) Overview of structure of baicalein-bound SARS-CoV-2 3CLpro. Protein is shown in cartoon representation and three domains are shown by different colors. Baicalein is shown as spheres with carbons in green. (**B**) 2*F*_o_-*F*_c_ density map contoured at 1.0 σ is shown for baicalein in the complex with SARS-CoV-2 3CLpro. The protein is colored gray. The substrate-binding pocket is represented by an inter-molecular surface. The inhibitor, baicalein, is shown as green sticks and a buried water molecular is displayed as a red ball. (**C**) Interactions formed between baicalein (green) and surrounding residues (cyan). Residues as well as the ligand are shown as sticks and hydrogen bonds are represented by black dashed lines.

## Crystal structure of SARS-CoV-2 3CLpro in complex with baicalein

The mode of action of baicalein and the structural determinants associated with its binding with SARS-CoV-2 3CLpro were further explored using X-ray protein crystallography. The crystal structure of SARS-CoV-2 3CLpro in complex with baicalein was determined at a resolution of 2.2 Å (Fig. 2; Table S2). The protease has a catalytic Cys145-His41 dyad and an extended binding site, features shared by SARS-CoV 3CLpro and MERS-CoV 3CLpro (Fig. 2A). The inhibitor binds in a core region of the substrate-binding site at the surface of the protease, between domains I and II (Fig. 2, A and B). Examination of the active site of the complex revealed that three phenolic hydroxyl groups of baicalein made multiple hydrogen bonds with main chains of Leu141/Gly143 as well as side chains of Ser144/His163 with or without the aid of a buried water molecule (Fig. 2C). The only carbonyl group established a hydrogen bond with the main chain of Glu166, while the free phenyl ring inserted into the S2 sub-pocket by making hydrophobic interactions with multiple residues Gln189/Arg188/Met49/Cys44/His41. Notably, apart from the hydrophobic interactions, the catalytic Cys145 and His41 also formed S—π (a distance of 3.3 Å from the sulfur to the centroid of the phenyl ring) and π—π interactions with the aromatic rings of baicalein, respectively. The side chain of Asn142 also established NH2—π (a distance of 3.6 Å from the nitrogen to the centroid of the phenyl ring) interactions with baicalein, which together with the interactions contributed by Cys145 lead the phenyl ring with three OH groups to be sandwiched between Cys145 and Asn142. Moreover, Met165 contacted the middle ring of baicalein via hydrophobic interactions. Consequently, baicalein is perfectly ensconced in the core of the substrate-binding pocket and interacts with two catalytic residues, the oxyanion loop (residues 138–145), Glu166, and the S1/S2 subsites, which are the key elements for recognition of substrates as well as peptidomimetic inhibitors (*17*). Although baicalein did not move deeply into the S1 sub-pocket, its phenolic hydroxyl groups did form contacts with the crucial residue of this sub-pocket, His163, via the water molecule. By the aid of an array of direct and indirect hydrogen bonds with Leu141/Gly141/Ser144, baicalein fixed the conformation of the flexible oxyanion loop, which serves to stabilize the tetrahedral transition state of the proteolytic reaction. These results together provide the molecular details of baicalein recognition by SARS-CoV-2 3CLpro and an explanation for the observed potent activity of such a small molecule against the protease.

## Unique binding mode of baicalein with SARS-CoV-2 3CLpro

The amino sequence of SARS-CoV-2 3CLpro displays 96% sequence identity to SARS-CoV 3CLpro. There are 12 residues differed in two proteases and none of them participates in the direct contacts with baicalein. The high level of sequence identified between two proteases allows one to assume that baicalein will bind to the SARS-CoV 3CLpro in the same way as it does to SARS-CoV-2 3CLpro. The inhibition assay shows that baicalein can also inhibit SARS-CoV 3CLpro, with an IC_50_ of 1.18 ± 0.37 μM. Thus, a three-dimensional model of SARS-CoV 3CLpro in complex with baicalein was constructed by superimposing the crystal structure of SARS-CoV-2 3CLpro/baicalein with that of SARS-CoV 3CLpro/TG-0204998 (PDB code 2ZU4) (Fig. S4A). The binding mode of baicalein with SARS-CoV 3CLpro is distinctively different from those of known inhibitors. All of the crystal structures of the inhibitor-bound SARS-CoV 3CLpro were collected for a comparison analysis (Fig. S4B). If those peptidomimetic inhibitors are delineated like “swords” to compete with the binding of substrate, baicalein works as a “shield” in front of two catalytic dyads to prevent the approach of the substrate to the active site (Fig. S4B). Such a unique binding mode in combination with its high ligand-binding efficiency and small molecular weight renders baicalein valuable for drug development.

## Antiviral activities of baicalin and baicalein in cells

We further evaluated the antiviral efficacy of baicalin and baicalein against a clinical isolate of SARS-CoV-2 (*5*) in Vero E6 cells. The cytotoxicity of two compounds in the cells was first determined by the CCK8 assay, and the resulting half-cytotoxic concentration (CC_50_) of the two compounds was over 200 μM, demonstrating a very low cytotoxicity of baicalin and baicalein (Fig. 3). Subsequently, the Vero E6 cells were infected with SARS-CoV-2 at a multiplicity of infection (MOI) of 0.05 in the presence of different concentrations of baicalin or baicalein. The antiviral efficacies were evaluated by quantification of viral copy numbers in the cell supernatant via quantitative real-time RT-PCR (qRT-PCR). As shown in Fig. 3, both baicalin and baicalein showed a dose-dependent inhibition on the replication of SARS-CoV-2, and the resulting half-maximal effective concentrations (EC_50_) of the two compounds were 10.27 and 1.69 μM, respectively. The corresponding selectivity index (SI = CC_50_/EC_50_) values are >19 and >118 for baicalin and baicalein, respectively. Therefore, the cell-based antiviral activity of baicalin or baicalein is superior to most of the reported compounds and that of baicalein is close to those of chloroquine (EC_50_: 1.13 μM; SI > 88) and remdesivir (EC_50_: 0.77 μM; SI > 129) (*18*).

**Fig. 3.**
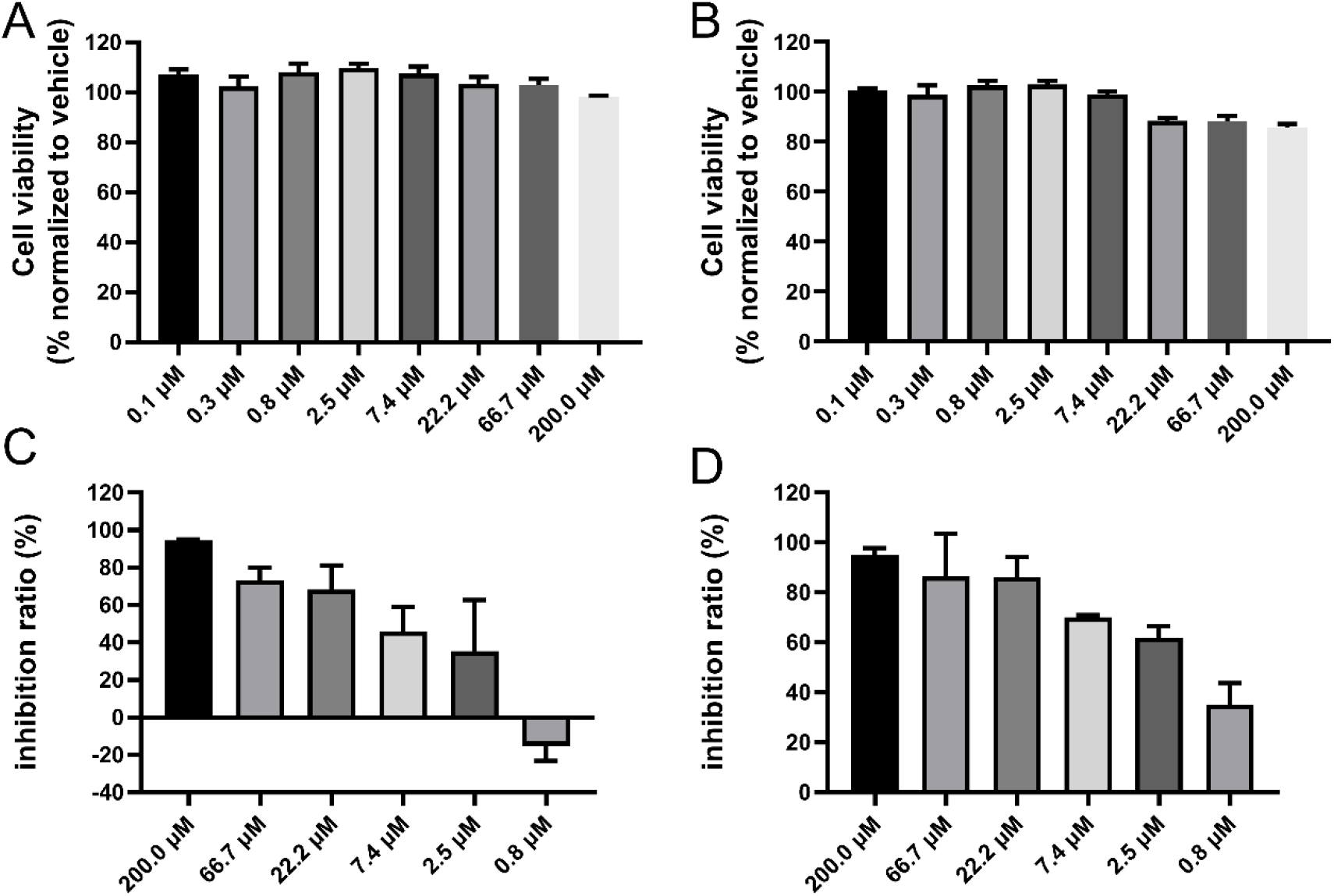
Antiviral activities of baicalin and baicalein against SARS-CoV-2 in Vero E6 cells. (**A** and **B**) Cytotoxicity of baicalin (A) or baicalein (B) to Vero E6 cells was measured by CCK-8 assay. (**C** and **D**) Cells were infected with SARS-CoV-2 at a multiplicity of infection (MOI) of 0.05 in the treatment of different concentrations of baicalin (C) or baicalein (D) for 48 h. Viral yield in the cell supernatant was then quantified by qRT-PCR.

## Discussion

Three global pandemics, SARS, MERS, and COVID-19, have raised great awareness about the increasing risks of highly pathogenic CoV infections in humans or animals and call for the development of efficacious anti-coronaviral drugs. 3CLpros provide a promising target for development of broad-spectrum antiviral drugs. These cysteine proteases are excised from the viral polyproteins by their own proteolytic activities and are indispensable for replication of CoVs. The substrate specificity of 3CLpros is featured by the efficient cleavage in the peptides including (Leu,Phe,Met,Val)-Gln↓(Ser,Ala,Gly) sequences (the cleavage site is indicated by ↓), and the crystal structures have revealed a remarkable degree of conservation of the substrate-binding sites, particularly for the crucial S1/S2 subsites (*7*, *19*, *20*). Accordingly, substrate analogs or mimetics attached with a chemical warhead targeting the catalytic cysteine were designed as peptidomimetic inhibitors of 3CLpros with a covalent mechanism of action (*6*). A series of diamide acetamides acting as non-covalent SARS-CoV 3CLpro inhibitors and their binding modes examined by crystal structures have been reported, but they are more or less peptidomimetic inhibitors and a continuous development of these compounds is absent (*21*). Although several other small molecules have been declared as 3CLpro inhibitors, a solid validation of their binding with 3CLpros by ITC or complex structure determination is lacking. As none of the known inhibitors has been moved to clinical trials, considerable efforts to discover novel small molecule inhibitors of 3CLpros are urgently needed.

In view of the long history of TCMs in treating viral infections and the urgent need of drug development against COVID-19, we investigated the anti-SARS-CoV-2 potential of natural products isolated from *S. baicalensis*, a long-term used TCM. *S. baicalensis* is also the major component of Chinese traditional patent medicines such as “Shuanghuanglian” and “Qingfei Paidutang” which have proven effective in curing patients of COVID-19 in China (*22*). As a result, we identified two small natural products, baicalin and baicalein, as the first class of non-covalent, non-peptidomimetic inhibitors of SARS-CoV-2 3CLpro by the enzymatic assay in combination with the ITC, ESI-MS, and X-ray protein crystallography studies. In contrast to most available covalent, peptidomimetic inhibitors of the 3CL proteases, the ITC measurement also revealed that these two compounds are non-covalent inhibitors with a high ligand-binding efficiency. The binding mode of baicalein with the protease revealed by the crystal structure demonstrates that a unique protein-ligand interaction pattern is utilized by baicalein to block the proteolytic activity of SARS-CoV-2 3CLpro. The crystal structure of baicalein-bound SARS-CoV-2 3CLpro thus reported the first demonstration, to the best of our knowledge, of the 3CLpro reversibly inhibited by a small molecule derived from TCM. With a molecular weight of 270.24 Da and a *K*_d_ of 4.03 μM, baicalein serves as the most efficient binder of 3CLpros known to date. Moreover, it possesses an antiviral activity in the cells with a low cytotoxicity. Given the high conservation of 3CLpro among various CoVs, baicalein is expected to inhibit other 3CL proteases of CoVs, providing a new candidate for development of broad-spectrum antiviral drugs.

In addition, baicalin tablets have been used as an adjuvant therapy for the treatment of acute, chronic, or persistent hepatitis in China. Single oral doses of 100–2800 mg of baicalein were safe and well tolerated by healthy subjects (clinical trials registration number CTR20132944) (*23*). Furthermore, a randomized, double-blind, placebo-controlled, multicenter, and phase IIa clinical trial for the effectiveness and safety of baicalein tablets in the treatment of improving other aspects of healthy adults with influenza fever is under way (clinical trials registration numbers CTR20182427 and NCT03830684). These favorable safety data together with the potent inhibitory activities in enzyme and cell-based assays show a strong preference for *in vivo* and clinical trials studies of baicalin and baicalein for COVID-19 treatment. Overall, the present study provides a good example for exploring the *in vitro* potency of TCMs and effectively identifying bioactive ingredients toward a specific target, especially when Chinese traditional patent medicines have been proven effective in curing patients of COVID-19 in China.

## Acknowledgments

The authors sincerely thank Prof. Zihe Rao and Prof. Haitao Yang for kindly providing the protein as well as the substrate for the enzymatic assay. We also thank the staff from beamlines BL17U1 and BL18U1 at Shanghai Synchrotron Radiation Facility. We thank LetPub (www.letpub.com) for its linguistic assistance during the preparation of this manuscript.

## Author contributions

H.S. and H.X. prepared the protein sample. H.S., W.Z., and H.X. performed the enzymatic assay and the ITC measurement. H.S., W.Z., and M.L. determined the crystal structure. S.Y., J. L., and C.K. parepared the compounds and carried out the mass spectroscopy experiments. W.S, Y.W., and L.Z. performed antiviral activities measurement in cells. M.G., K.Y., H.L., J.S., W.T., and J.Z. helped with data analysis and interpretation. Y.X., H.J., Y.Y., and F.B. initiated the project and supervised the research. Y.X. wrote the manuscript with input from all co-authors.

## Data availability statements

The SARS-CoV-2 3CLpro–baicalein complex structure was deposited with the Protein Data Bank with accession code 6M2N. All other data are available from the corresponding author upon reasonable request.

## Supplementary Materials

Materials and Methods

Figures S1–S4

Tables S1 and S2

References (24–29)

## Supplementary Materials

### Materials and Methods

#### General chemistry

*S. baicalensis* Georgi decoction pieces (No. 191226) were purchased from Beijing Tongrentang China Pharmacy in Shanghai and its aqueous extract was prepared in accordance with clinical application. The fractionation of *S. baicalensis* aqueous extract was performed on a Waters AutoPurification HPLC with a 2489 PDA detector coupled with a Waters Acquity ELSD using a Waters Sunfire RP C18 column (30 × 150 mm, 5 μm) with a 30.0 mL/min flow rate. Baicalin and baicalein were isolated from active fractions from *S. baicalensis* with a purity over 95%. Analytical HPLC and ESIMS spectra were performed on a Waters 2695 instrument with a 2998 PDA detector coupled with a Waters Acquity ELSD and a Waters 3100 SQDMS detector using a Waters Sunfire RP C18 column (4.6 × 150 mm, 5 μm) with a 1.0 mL/min flow rate. ^1^H and ^13^C NMR spectra were recorded on a Bruker AVANCE III 600 MHz instrument. Chemical shifts were reported in ppm (*δ*) coupling constants (*J*) in hertz. Chemical shifts are reported in ppm units with Me_4_Si as a reference standard.

#### Protein expression and purification

The cDNA of full length SARS-CoV-2 3CLpro or SARS-CoV 3CLpro was cloned into the PGEX6p-1 vector. To obtain SARS-CoV-2 3CLpro or SARS-CoV 3CLpro with authentic N and C terminals, four amino acids (AVLQ) were inserted between the GST tag and the full length SARS-CoV-2 3CLpro or SARS-CoV 3CLpro, while eight amino acid (GPHHHHHH) were added to the C-terminal of SARS-CoV-2-3CLpro. The plasmid was then transformed into BL21 (DE3) cells for protein expression. The N terminal GST tag and four amino acids (AVLQ) was self-cleavable. The expressed protein with authentic N terminal was purified by a Ni-NTA column (GE Healthcare) and transformed into the cleavage buffer (150 mM NaCl, 25 mM Tris, pH 7.5) containing human rhinovirus 3C protease for removing the additional residues. The resulting protein sample was further passed through a size exclusion chromatography (Superdex200, GE Healthcare). The eluted protein samples were stored in a solution (10 mM Tris, pH 7.5) for the enzymatic inhibition assay, native state mass spectrometry studies and protein crystallization.

To obtain SARS-CoV-2-3CLpro with the GST tag, the cDNA of full length SARS-CoV-2-3CLpro was inserted after the prescission protease digestion site of the PGEX6p-1 vector. The plasmid was transformed into BL21 (DE3) cells for protein expression. The expressed protein was purified by a GST column (GE Healthcare) followed by the size exclusion chromatography (Superdex200, GE Healthcare). The resulting protein was stored in a solution (25 mM Tris, pH 7.3) for the isothermal titration calorimetry measurements.

#### The inhibition assay of SARS-CoV-2-3CLpro

A fluorescence resonance energy transfer (FRET) protease assay was applied to measure the inhibitory activity of compounds against the SARS-CoV-2 3CLpro or SARS-CoV 3CLpro. The fluorogenic substrate (MCA-AVLQSGFR-Lys(Dnp)-Lys-NH2) was synthesized by GenScript (Nanjing, China). The FRET-based protease assay was performed as follows. The recombinant SARS-CoV-2-3CLpro (30 nM at a final concentration) or SARS-CoV 3CLpro (100 nM at a final concentration) was mixed with serial dilutions of each compound, oral liquid or the dissolved lyophilized powder in 80 μL assay buffer (50 mM Tris–HCl, pH 7.3, 1 mM EDTA) and incubated for 10 min. The reaction was initiated by adding 40 μL fluorogenic substrate with a final concentration of 20 μM. After that, the fluorescence signal at 320 nm (excitation)/405 nm (emission) was immediately measured every 30 s for 10 min with a Bio-Tek Synergy4 plate reader. The *Vmax* of reactions added with compounds at various concentrations compared to the reaction added with DMSO were calculated and used to generate IC_50_ curves. For each compound, at least three independent experiments were performed for the determination of IC_50_ values and each experiment was carried out in triplicates. The IC_50_ values were expressed as mean ± SD from three independent experiments and determined via the nonlinear regression analysis using GraphPad Prism software 8.0 (GraphPad Software, Inc., San Diego, CA, USA).

#### Separation of *S. baicalensis* extract

*S. baicalensis* decoction pieces (75 g) were boiled in 750 ml of water for 1 hour twice. After filtration, the combined aqueous extract was evaporated under reduced pressure and the residue was then dried by lyophilization. 455 mg of the dried extract was subjected to Waters AutoPurification HPLC eluting with 5% ~ 60% MeCN in water containing 0.1% HCOOH from 0-40min. All of the eluent fractions were collected by automatic fraction collector and combined by peaks on UV (280 and 320 nm) and ELSD detectors. Totally 14 fractions were obtained with a recovery rate of 92% for the assay of SARS-CoV-2-3CLpro inhibition.

#### Isothermal titration calorimetry (ITC) measurements

All measurements were performed by iTC200 calorimeter (General Electric Co.) in an ITC buffer (25 mM Tris, pH 7.3) while stirring at 800 rpm. The stock solution of compounds and the SARS-CoV-2-3CLpro protein with a GST tag were diluted with the ITC buffer to a compound concentration of 50-100 μM and a protein concentration of 0.6-1.5 mM before titrations. The final concentration of DMSO in the reaction buffer is less than 2% of the total volume. Compounds were titrated by the protein solution. All titrations were performed using an initial injection of 0.4 μL followed by 19 identical injections of 2 μL with a duration of 4 seconds per injection and a spacing of 120 seconds between injections. The last five data points were averaged and subtracted from each titration to account for the heat of dilution. Additional background experiments where the protein solution was titrated into the buffer without the compound revealed no significant shift in the baseline during the course of the measurements. Each measurement was repeated three times.

#### Native state mass spectrometry studies

All mass spectrometry (MS) measurements were performed with a scimaX MRMS system (Bruker Daltonik GmbH, Bremen, Germany). The sample solutions were infused to the system with syringe (50 μL) at a flow rate of 120 μL/h using electrospray ionization in a positive ion mode. Data were acquired in the mass-to-charge (*m/z*) range 400-6000 with a transient length of 3.92 s in quadrupolar detection resulting in a resolving power of 54.0 ppb at m/z 3240. The source and ion transfer parameters were optimized as follows: Skimmer 1 100 V, Funnel 1 180 V, Funnel RF Amplitude 200 V, collision cell frequency 2.0 MHz, ion transfer frequency 2 MHz, time of flight to analyzer cell 2.0 ms with RF amplitude of 350 V, and source temperature 200°C. For the protein performance optimization, the mass range around the change stated 18+ was isolated with the quadruple using a quadrupole mass window of 200 Da with a center mass of the quadrupole of *m/z* 3750; the magnitude size was set to 4M; 200 single scans were added. For ligand binding screening and dissociation constant (K_d_) determination, the magnitude size was set to 128 K and 1000 single scans were added.

The protein sample of SARS-CoV-2-3CLpro was buffer exchanged with 10 mM ammonium acetate buffer (pH 6.8). For optimization of the MS condition, the protein was diluted into 2 μM. For ligand binding screening studies and *K*_d_ determination, samples was prepared with the protein at 1 μM and ligands at a series of concentrations (50, 25, 12.5, 6.25, 3.13, 1.56, 0.78 μM for baicalin and 10, 5, 2.5, 1.25, 0.63, 0.31, 0.16, 0.08 μM for baicalein). Incubation time was 1 h at 37°C. Three independent experiments were performed for the determination of *K*_d_s.

Mass spectra were deconvoluted with SNAP2 in DataAnalysis 5.2 (Bruker Daltonics, Bremen, Germany). The charge states 18+ and 19+ have been used for calculation of the free protein and protein-ligand complex intensities. The relative abundances of protein-ligand complex to total protein in the mass spectra were calculated as reported previously (*24*).

#### Protein crystallization and structure determination

The purified SARS-CoV-2-3CLpro protein was concentrated to 7 mg/mL for crystallization. One hour incubation of the protein with 10 mM baicalein was carried out before crystallization condition screening. Crystals of the complex were obtained at 20 °C by mixing equal volumes of protein-baicalein and a reservoir (16 % PEG6000, 100 mM MES, pH 5.8, 3% DMSO) with a handing-drop vapor diffusion method. Crystals were flash frozen in liquid nitrogen in the presence of the reservoir solution supplemented with 20% glycerol. X-ray diffraction data were collected at beamline BL18U1 at the Shanghai Synchrotron Radiation Facility (*25*). The data were processed with HKL3000 software packages (*26*). The complex structure was solved by molecular replacement using the program PHASER (*27*) with a search model of PDB code 6LU7. The model was built using Coot (*28*) and refined with a simulated-annealing protocol implemented in the program PHENIX (*29*). The refined structure was deposited to Protein Data Bank with an accession code listed in Table S1. The complete statistics as well as the quality of the solved structure are also shown in Table S1.

#### The cell-based antiviral activity assay

The Vero E6 cell line was obtained from American Type Culture Collection (ATCC, Manassas, USA) and maintained in minimum Eagle’s medium (MEM; Gibco Invitrogen) supplemented with 10% fetal bovine serum (FBS; Invitrogen, UK) in a humid incubator with 5% CO2 at 37 °C. The cytotoxicity of two tested compounds on the Vero E6 cells were determined by CCK8 assays (Beyotime, China). A clinical isolate SARS-CoV-2 (*5*) was propagated in the Vero E6 cells, and the viral titer was determined as described previously (*18*). All the infection experiments were performed at biosafety level-3 (BLS-3).

Pre-seeded Vero E6 cells (5×10^4^ cells/well) were incubated with different concentrations of baicalin or baicalein for 1 hour and the virus was subsequently added (a multiplicity of infection of 0.05) to infect the cells for 2 hours. After that, the virus-compound mixture was removed and cells were further cultured with a fresh compound containing medium. At 48 h p.i., the cell supernatant was collected and the viral RNA in supernatant was subjected to qRT-PCR analysis as described previously (*18*). DMSO was used in the controls. The experiments were performed in triplicates and three independent experiments were carried out for each compound.

**Fig. S1.**
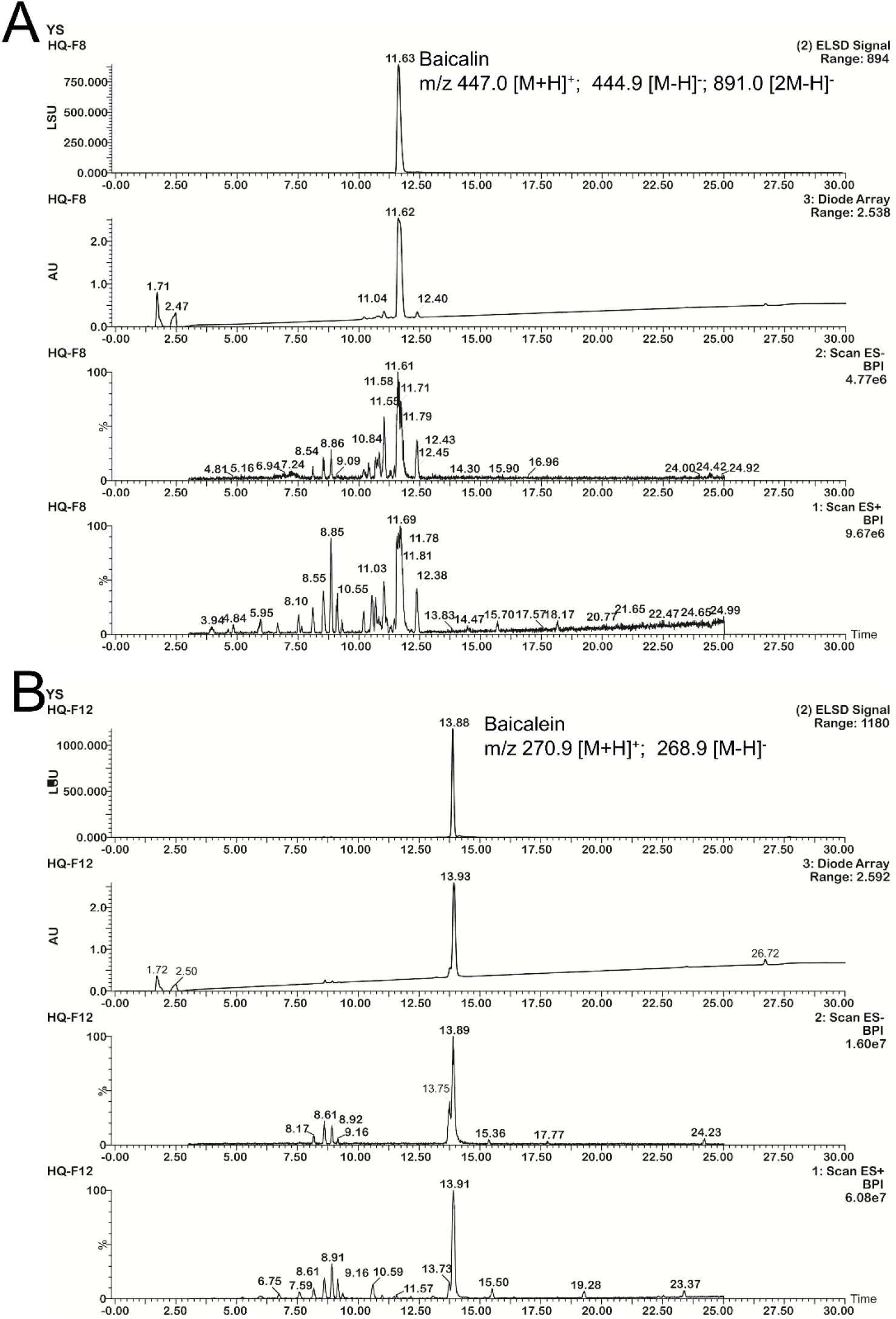
HPLC-MS profiling of the active fraction 8 and 12. (**A**) HPLC-MS profiling of fration 8. (**B**) HPLC-MS profiling of fration 12

**Fig. S2.**
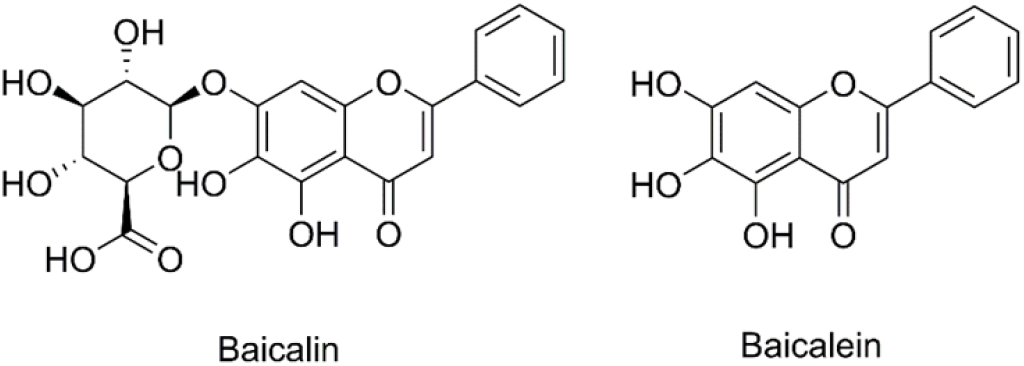
Chemical structures of baicalin, and baicalein.

**Fig. S3.**
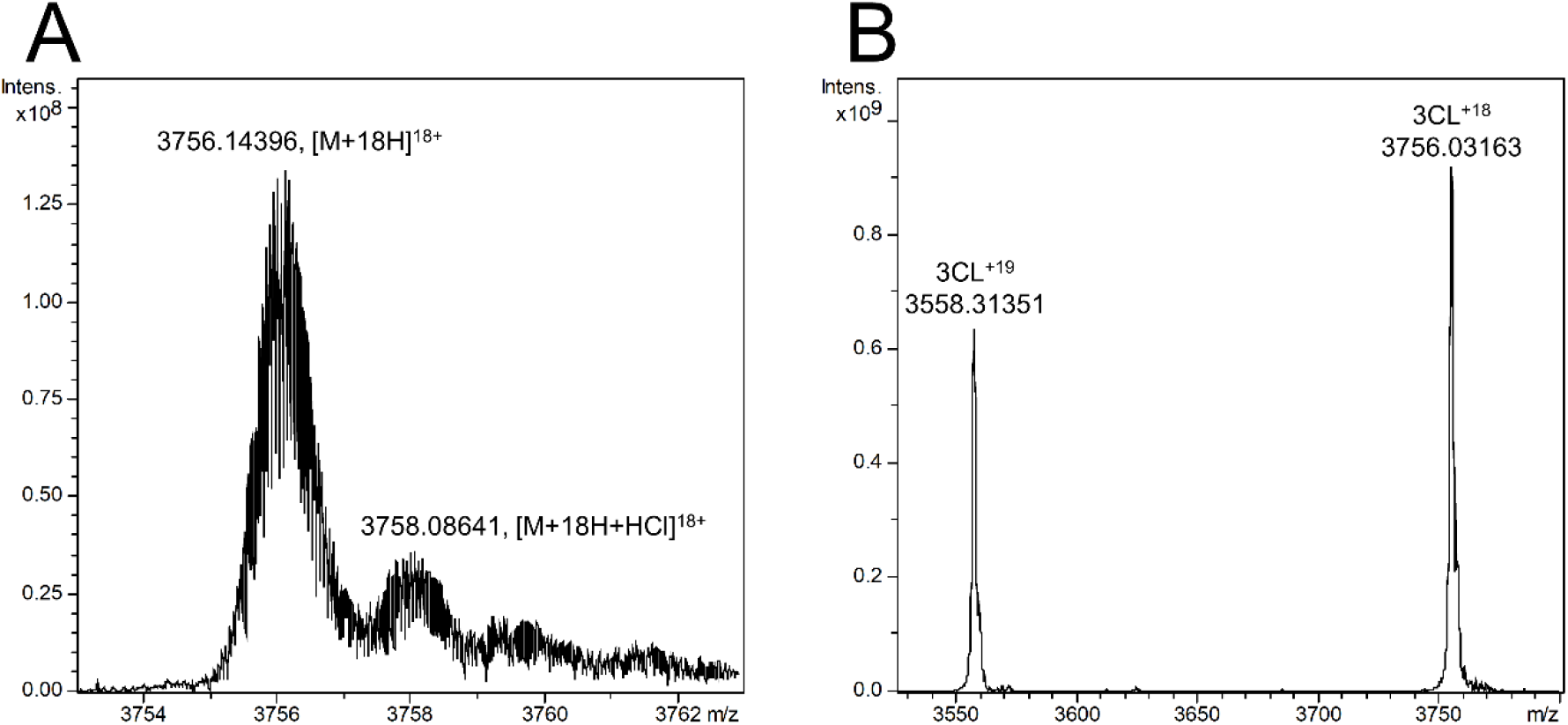
The native state mass spectrometry spectro of SARS-CoV-2-3CLpro. (**A**) The 18+ charge state of SARS-CoV-2-3CLpro under an optimization condition. (**B**) SARS-CoV-2-3CLpro under a protein-ligand binding screening condition.

**Fig. S4.**
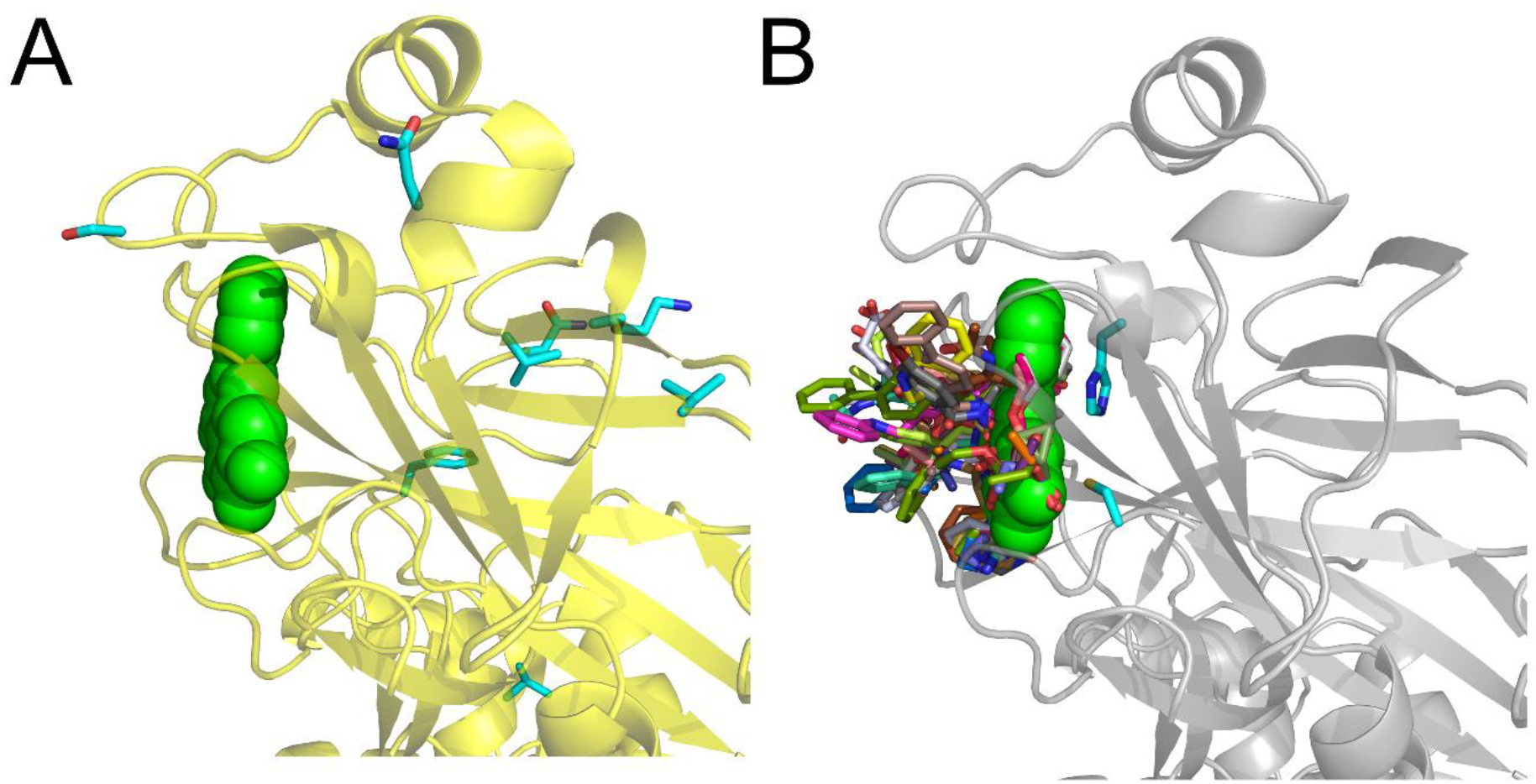
The unique binding mode of baicalein with SARS-CoV-3CLpro. (**A**) The structure of SARS-CoV-3CLpro in complex with baicalein was generated by superimposing the structure of SARS-CoV-2-3CLpro/baicalein (PDB code 6M2N) with that of SARS-CoV-3CLpro/TG-0204998 (PDB code 2ZU4). The SARS-CoV-3CLpro is shown in yellow cartoon and baicalein is shown as spheres with carbons in green. Residues differed in two proteases are shown as cyan sticks. (**B**) A survey of the binding modes of known inhibitors as well as baicalein with SARS-CoV-3CLpro are shown by superimposing all the crystal structures of inhibitor-bound SARS-CoV-3CLpro (PDB codes: 2A5I, 2A5K, 2ALV, 2AMD, 2AMQ, 2GTB, 2GX4, 2OP9, 2YNB, 2ZU4, 2ZU5, 3ATW, 3AVZ, 3AW0, 3SN8, 3SNA, 3SNB, 3SNC, 3SND, 3SNE, 3SZN, 3V3M, 4MDS, 4TWW, 4TWY, 4WY3, 5C5N, and 5C5O). Baicalein is shown as green spheres and other inhibitors together with two catalytic residues are shown as sticks.

**Table S1.** Inhibition of each fraction from the aqueous extract of *S. baicalensis* on SARS-CoV-2-3CLpro

| code | Inhibition ratio (%) |  |  |  | Major component(s) |
| --- | --- | --- | --- | --- | --- |
|  | 10.0 µg/mL | 2.0 µg/mL | 0.4 µg/mL | 0.08 µg/mL |  |
| F1 | 6.4 |  |  |  |  |
| F2 | 17.9 |  |  |  |  |
| F3 | -0.6 |  |  |  |  |
| F4 | 6.2 |  |  |  |  |
| F5 | 6.9 |  |  |  |  |
| F6 | 15.1 |  |  |  |  |
| F7 | 28.3 |  |  |  |  |
| F8 | 60.3 | 19.0 |  |  | Baicalin |
| F9 | 7.4 |  |  |  |  |
| F10 | 6.2 |  |  |  |  |
| F11 | 102.3 | 94.2 | 57.9 | 21.9 | Baicalein |
| F12 | 19.8 |  |  |  |  |
| F13 | 3.8 |  |  |  |  |
| F14 | 3.2 |  |  |  |  |
| Total Extract | 68.6 | 18.3 |  |  |  |

**Table S2.**
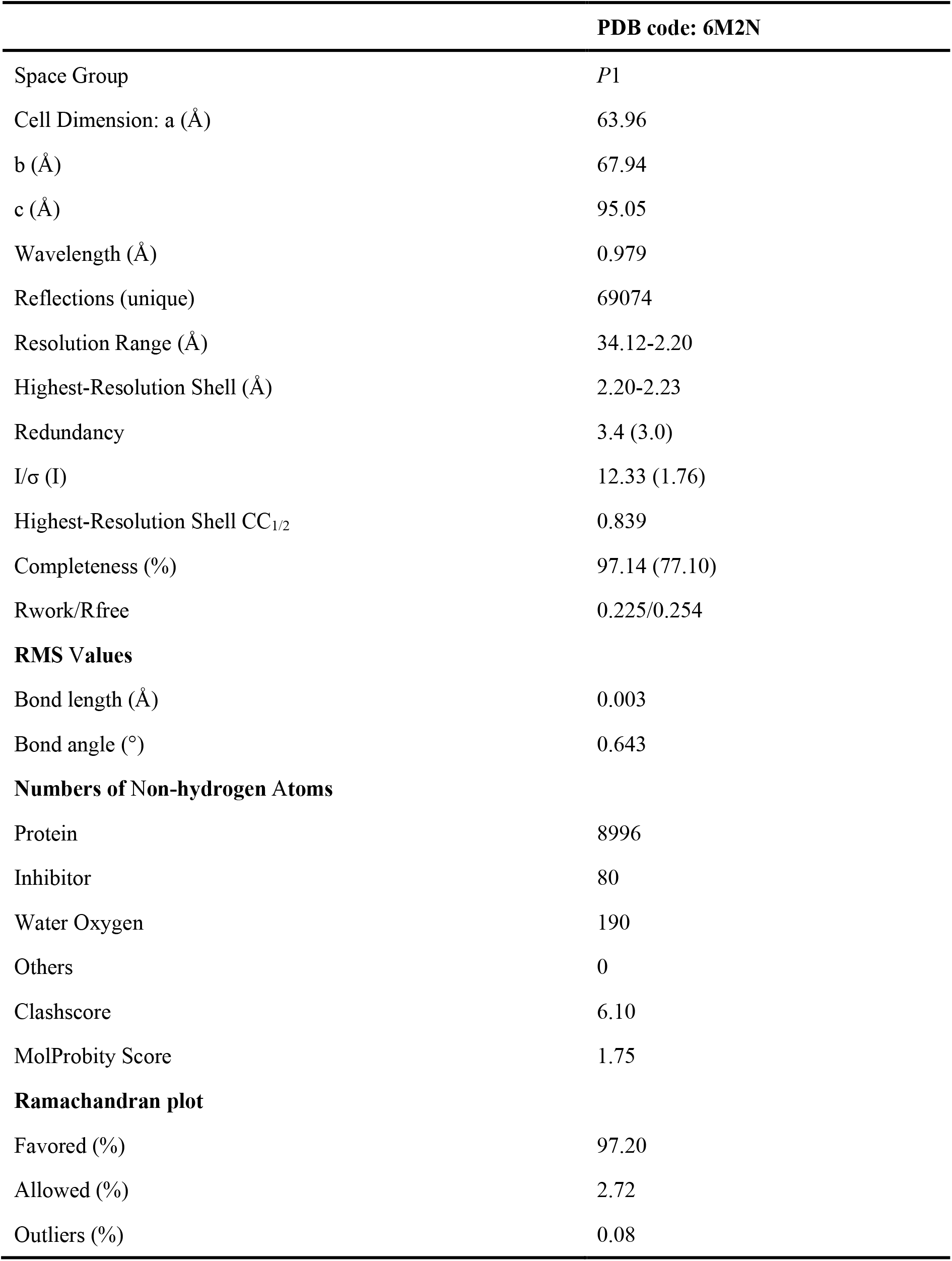
Crystallography data collection and refinement statistics.

